# DNA repair is essential for *Vibrio cholerae* growth on Thiosulfate-Citrate-Bile Salts-Sucrose (TCBS) Medium

**DOI:** 10.1101/2025.01.10.632459

**Authors:** Alex J. Wessel, Drew T.T. Johnson, Christopher M. Waters

## Abstract

Thiosulfate-citrate-bile salts-sucrose (TCBS) agar is a selective and differential media for the enrichment of pathogenic *Vibrios*. We observed that an exonuclease VII (*exoVII*) mutant of *Vibrio cholerae* failed to grow on TCBS agar, suggesting that DNA repair mutant strains may be hampered for growth in this selective media. Examination of the selective components of TCBS revealed that bile acids were primarily responsible for toxicity of the *exoVII* mutant. Suppressor mutations in DNA gyrase restored growth of the *exoVII* mutants on TCBS, suggesting that TCBS inhibits DNA gyrase similar to the antibiotic ciprofloxacin. To better understand what factors are important for *V. cholerae* to grow on TCBS, we generated a randomly-barcoded TnSeq (RB-TnSeq) library in *V. cholerae* and have used it to uncover a range of DNA repair mutants that also fail to grow on TCBS agar. The results of this study suggest that TCBS agar causes DNA damage to *V. cholerae* similarly to the mechanism of action of fluoroquinolones, and overcoming this DNA damage is critical for *Vibrio* growth on this selective medium.

**Abstract Importance:** TCBS is often used to diagnose cholera infection. We found that many mutant *V. cholerae* strains are attenuated for growth on TCBS agar, meaning they could remain undetected using this culture-dependent method. Hypermutator strains with defects in DNA repair pathways might be especially inhibited by TCBS. In addition, *V. cholerae* grown successively on TCBS agar develops resistance to ciprofloxacin.

## Introduction

*Vibrio cholerae* is the causative agent of cholera, a gastrointestinal infection characterized by profuse watery diarrhea that, when left untreated, causes rapid dehydration and death. This gram-negative bacterium lives a dual lifestyle, alternating between its aquatic environmental niche and human host environment where it is acquired through ingestion of contaminated food or water. Cholera remains endemic in much of the developing world, with sporadic outbreaks occurring in nations with poor sanitation practices and limited access to clean drinking water (1, 2).

Culture confirmation from stool samples remains the gold standard for the diagnosis of *V. cholerae* infections (3). This is typically achieved using Thiosulfate-Citrate-Bile Salts-Sucrose (TCBS) agar, a highly selective medium for *Vibrios*. Following growth, the identity of a *Vibrio* grown on TCBS agar can be further determined based on its capacity to ferment sucrose— while *V. cholerae* can ferment sucrose, most other medically relevant *Vibrios* cannot (4).

Aside from sucrose, the major selective factors in TCBS are the medium’s alkaline pH and bile salts, which inhibit the growth of non-*Vibrio* enterics. Given that *V. cholerae* encounters bile routinely throughout its pathogenic lifecycle, it has evolved a variety of mechanisms to sense, respond to, and limit the antimicrobial effects of bile. To this end, bile represents a significant environmental signal in the pathogenesis of *V. cholerae*. *V. cholerae* senses bile through an interaction between the inner membrane sensory and regulatory proteins ToxR and ToxS (5, 6). For example, in the human intestinal tract, bile activation of ToxRS adapts *V. cholerae* for bile tolerance by inducing and inhibiting the expression of outer membrane proteins OmpU and OmpT, respectively (7–9). Because bile can enter the cell through OmpT but not OmpU (10, 11), a shift in porin expression protects *V. cholerae* from its cytotoxic effects.

The bacterial stress response to bile is mostly characterized by DNA damage leading to SOS induction as well as the remodeling of the cell membrane (12–17). Early works in *E. coli* demonstrate that cell death and SOS induction in response to bile mimic that of cells treated with the DNA crosslinking agent mitomycin C, suggesting that bile may cause direct DNA damage in bacteria (18). Another study by Prieto and colleagues indicate that treatment of *Salmonella enterica* with bile causes oxidative DNA damage and increases the frequency of GC → AT transitions. This led the authors to suggest that base excision repair and recombinational repair, but not nucleotide excision repair, are necessary for *S. enterica* tolerance to bile (13).

In this work we demonstrate that exonuclease VII (*exoVII*) mutant *V. cholerae* fails to grow on TCBS agar, and that the main component in TCBS agar that is responsible for this inhibition is ox bile. We found that mutations in DNA gyrase (encoded by genes *gyrA* and *gyrB*) suppress the toxicity of TCBS in the absence of ExoVII and that some of these mutations confer resistance to the fluoroquinolone ciprofloxacin. Finally, we performed randomly barcoded transposon insertion site sequencing (RB-TnSeq) to identify other *V. cholerae* mutants with growth defects on TCBS agar, and in doing so uncovered mutants with component-dependent growth defects. The results presented here (19) provide an explanation for the inhibition of *exoVII* mutant *V. cholerae* and suggest that bile salts/acids may induce quinolone-like DNA damage in bacteria. Moreover, we show that DNA repair is essential for *V. cholerae* to robustly grow on TCBS which has significant implications on whether TCBS can accurately sample the diversity of *V. cholerae* in clinical isolates.

## Results

### *xseA* mutant *V. cholerae* cannot grow on TCBS agar

While conducting previously-described experiments to enhance the efficiency of Multiplex Genome Editing by Natural co-Transformation (MuGENT)(28), we serendipitously discovered that a *V. cholerae* mutant strain derived from the El Tor Biotype E7946 (strain TND0252: Ptac-*tfoX,* Δr*ecJ*501bp, Δ*exoVII*501bp, Δ*lacZ*::*lacIq*, Δ*vc1807*::SpecR), which had been developed to allow MuGENT with minimal homology regions, failed to grow on Thiosulfate-Citrate-Bile Salts-Sucrose (TCBS) agar, a selective and differential medium for *Vibrios*. However, the parental E7946 strain and another El Tor isolate that is widely used, C6706, grew well (Fig 1A). TND0252 has three notable changes including Isopropyl β-d-1-thiogalactopyranoside (IPTG)-inducible *tfoX*, which triggers natural competence, and null mutations in the two nucleases *recJ* and *exoVII. vc1807* is a frame-shifted transposase that appears to be neutral in all conditions and is used in the MuGENT protocol and thus likely was not responsible for the lack of growth on TCBS (28). We therefore hypothesized mutation of *tfoX, recJ,* or *exoVII* was responsible for TCBS toxicity. We found that repairing the *tfoX* (strain CW2171, “+P*tfox*”) or *recJ* gene (strain CW2173, “+*recJ*”) with WT sequence did not restore WT-like growth on TCBS agar. However, restoration of the *xseA* null mutation to wild type sequence (strain CW2172, “+*exoVII*”) restored TCBS growth (Fig. 1A). *xseA,* along with *xseB,* form the ExoVII nuclease that has been implicated in DNA repair. We thus conclude that *V. cholerae* mutants lacking functional ExoVII are highly attenuated for growth on TCBS agar.

**Figure 1:**
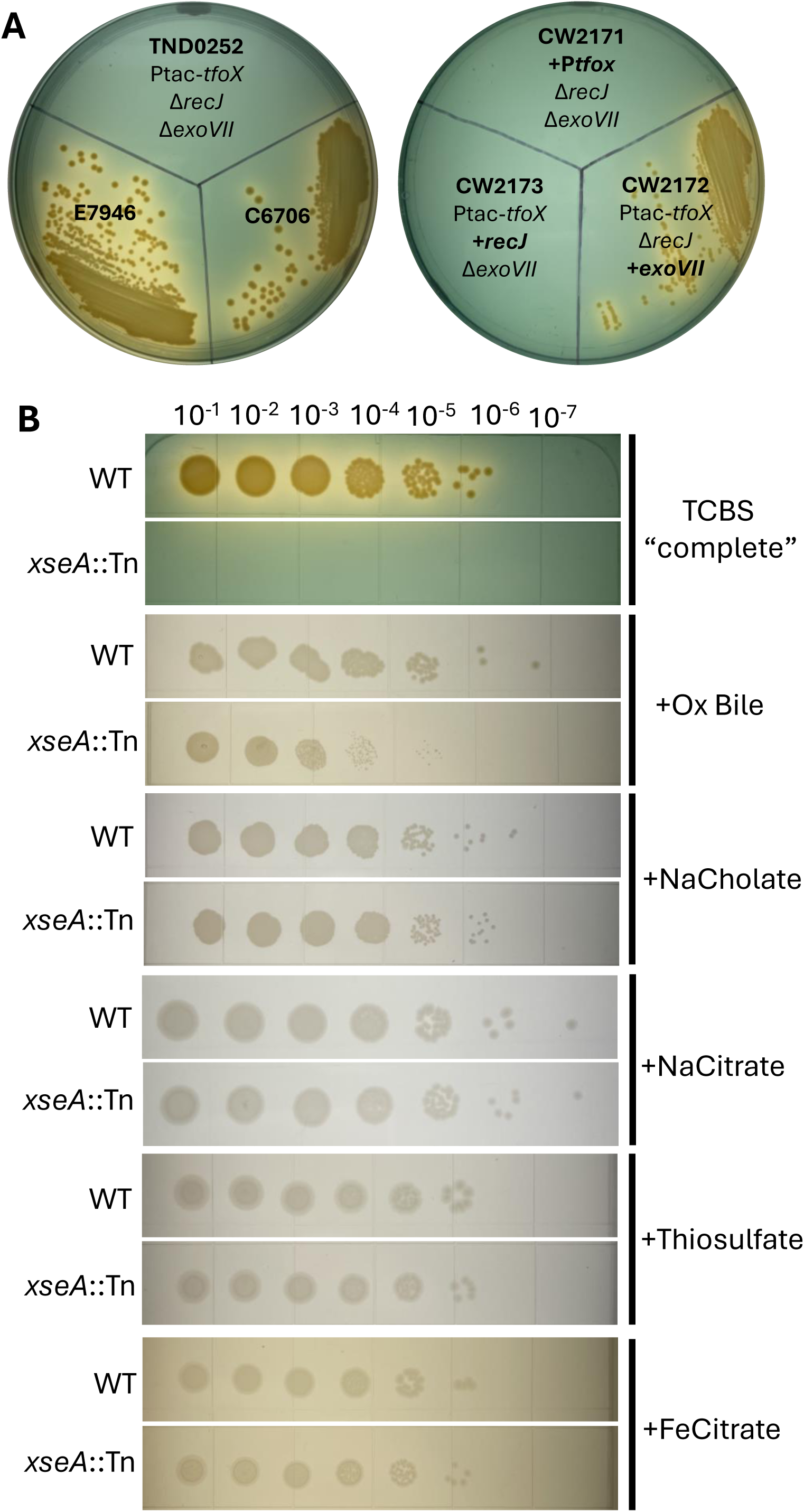
*exoVII* mutant *V. cholerae* cannot grow on TCBS agar. (A) Parent strains (left plate) and repaired strains (right plate) were struck over TCBS agar and assayed for growth after 24 hours incubation. Strain E7946 is the parent to TND0252. TND0252 is the parent strain to CW2171, CW2172 and CW2173. (B) Serial dilutions of WT and *xseA*::Tn *V. cholerae* spotted over TCBS agar, or TCBS base agar supplemented with a single selective component as indicated using the wt/vol listed in Table 1. For both figures A and B, plates shown are representative of 3 biological replicate experiments.

**Table 1.** Components of Thiosulfate-Citrate-Bile Salts-Sucrose medium. Individual components masses per 1L of TCBS broth. Components thought to be selective are written in bold. pH ∼8.6 at 25°C.

|  |  |
| --- | --- |
| Sucrose | 20g |
| Peptone | 10g |
| Yeast extract | 5g |
| Sodium chloride | 10g |
| **Sodium citrate** | 10g |
| **Ox bile** | 5g |
| **Sodium cholate** | 3g |
| **Ferric citrate** | 1g |
| **Sodium thiosulfate** | 10g |

We next questioned what component of TCBS agar was responsible for inhibiting the *exoVII* mutant. TCBS agar contains several selective components that inhibit the growth of non-*Vibrio* enteric bacteria (Table 1). We retrieved and cultured a mariner transposon *xseA* mutant (hereafter *xseA::*Tn) from a regenerated ordered *V. cholerae* strain C6706 mutant library (21) and spot plated serial dilutions of this mutant and the WT strain over different compositions of TCBS agar. As seen with strain E7946, the *xseA* null mutant of C6706 failed to grow on TCBS (Fig. 1B). We reasoned the non-selective components of TCBS include sucrose, peptone, yeast extract, and sodium chloride, and media containing only these four ingredients was termed “TCBS base” media. Alternatively, the other components of TCBS including sodium citrate, ox bile, sodium cholate, ferric citrate, and thiosulfate likely provide selection for *Vibrios*. To identify which selective component of TCBS inhibited growth of the *xseA*::Tn mutant, we spot plated this mutant and WT on base TCBS containing only one of the selective components. The *xseA::*Tn mutant grew equivalent to WT in TCBS base agar supplemented with sodium cholate, sodium citrate, sodium thiosulfate, or ferric citrate added at the wt/vol listed in Table 1, but was strongly attenuated relative to WT on TCBS base agar supplemented with ox bile (Fig. 1B). Of note, *xseA::*Tn did exhibit limited growth on ox bile. Thus, we concluded the major selective component responsible for inhibiting the *exoVII* mutant *V. cholerae* strain is ox bile although growth on complete TCBS was the most attenuated.

### *exoVII* growth inhibition on TCBS can be suppressed by mutations in DNA gyrase

To better understand the requirement for *exoVII* growth on TBCS, we plated the *xseA*::Tn strain on TCBS agar to select for suppressor mutations that restore growth. We isolated and sequenced the genomes of four suppressor mutants. Three of the four mutants had unique mutations in either of the two subunits of DNA gyrase- *gyrA* and *gyrB* (Fig. 2A, Table S1). Of note, one of the four mutants had mutations in both *gyrA* and gyrB (Fig. 2A, Table S1). As DNA gyrase is essential for growth (29–31), we deduce that these mutations did not produce a null phenotype but rather altered the properties of DNA gyrase restoring TCBS growth. The suppressor that did not encode a mutation in *gyrA* or *gyrB* mapped to the ATP-dependent DNA helicase *dinG* (GPY04_RS07870) (Table S1), but for this study we will focus on the more common DNA gyrase mutations. We used MuGENT to regenerate *the gyrA/gyrB* suppressor mutations in the parent *xseA*::Tn strain and confirmed that promote growth on TCBS agar despite the lack of functional ExoVII (Fig. 2B).

**Figure 2:**
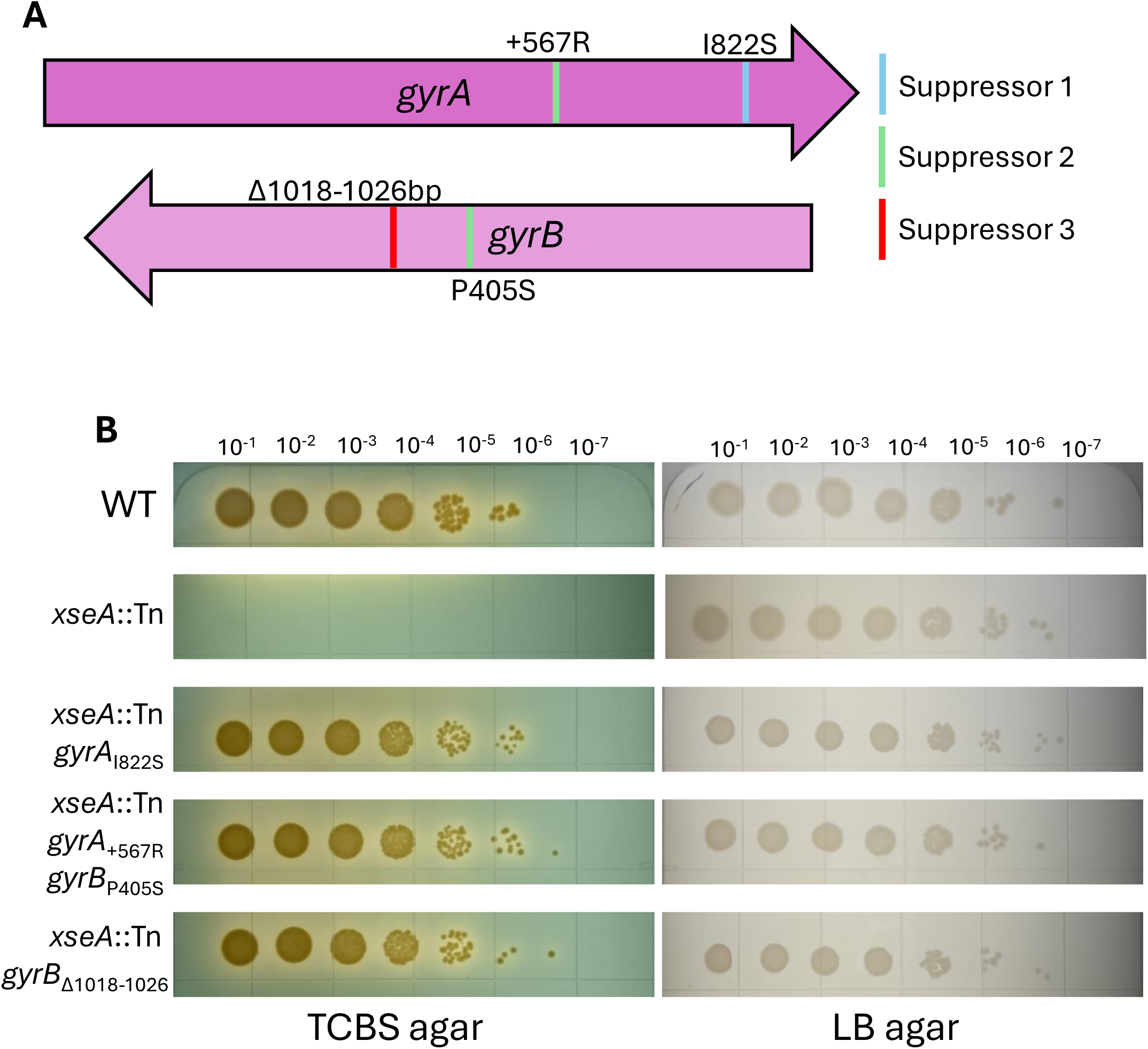
Inhibition of *exoVII* mutants on TCBS agar can be suppressed by mutations in DNA gyrase. (A) Together, *gyrA* and *gyrB* encode DNA gyrase. Color-coded bands within the *gyrA* and *gyrB* reading frames correspond to mutations present among three suppressor mutants generated by plating *xseA*::Tn over TCBS agar and selecting isolated colonies once growth was observed. (B) Serial dilution and spot plating of WT, *xseA*::Tn and *exoVII* suppressor mutants over TCBS agar and LB agar. Plates shown are representative of 3 biological replicate experiments.

### *exoVII* and *gyrAB* alter *V. cholerae* ciprofloxacin resistance

DNA gyrase, also known as bacterial type II topoisomerase, relieves topological strain on unwound DNA by performing negative supercoiling, and is the target of quinolone antibiotics. Quinolone drugs like ciprofloxacin work as topoisomerase poisons by trapping DNA gyrase in a covalently attached DNA complex after it makes double stranded breaks (DSBs), causing cell death by inhibiting DNA replication and transcription while leading to the accumulation of DSBs (32). In *E. coli,* ExoVII mutants are hypersensitive to ciprofloxacin, and thus ExoVII was hypothesized to relieve covalent complexes of DNA gyrase bound to DNA (19). To test whether ExoVII has a similar function in *V. cholerae*, we measured the sensitivity of ExoVII null mutants to ciprofloxacin using a disk diffusion assay. We found that, indeed, mutant *V. cholerae* lacking either subunit of *exoVII* (*xseA*::Tn or *xseB*::Tn) are indeed significantly more sensitive to inhibition by ciprofloxacin than WT exhibiting a larger zone of inhibition (ZOI) (Fig. 3A). We next hypothesized that suppressor mutations in *gyrA* and/or *gyrB* would reduce ciprofloxacin sensitivity in the *xseA*:Tn mutant. However, contrary to our prediction only one of the three suppressor mutant strains (*xseA*::Tn *gyrB*_Δ1018-1026_) significantly restored WT ciprofloxacin resistance whereas the other two maintained ciprofloxacin sensitivity similar to *xseA:*Tn (Fig. 3A). Therefore, even though all three DNA gyrase suppressor mutants were able to overcome *xseA*:Tn growth inhibition on TCBS, only one leads to enhanced ciprofloxacin resistance, suggesting that the disruption of DNA gyrase by TCBS is mechanistically distinct from ciprofloxacin.

**Figure 3:**
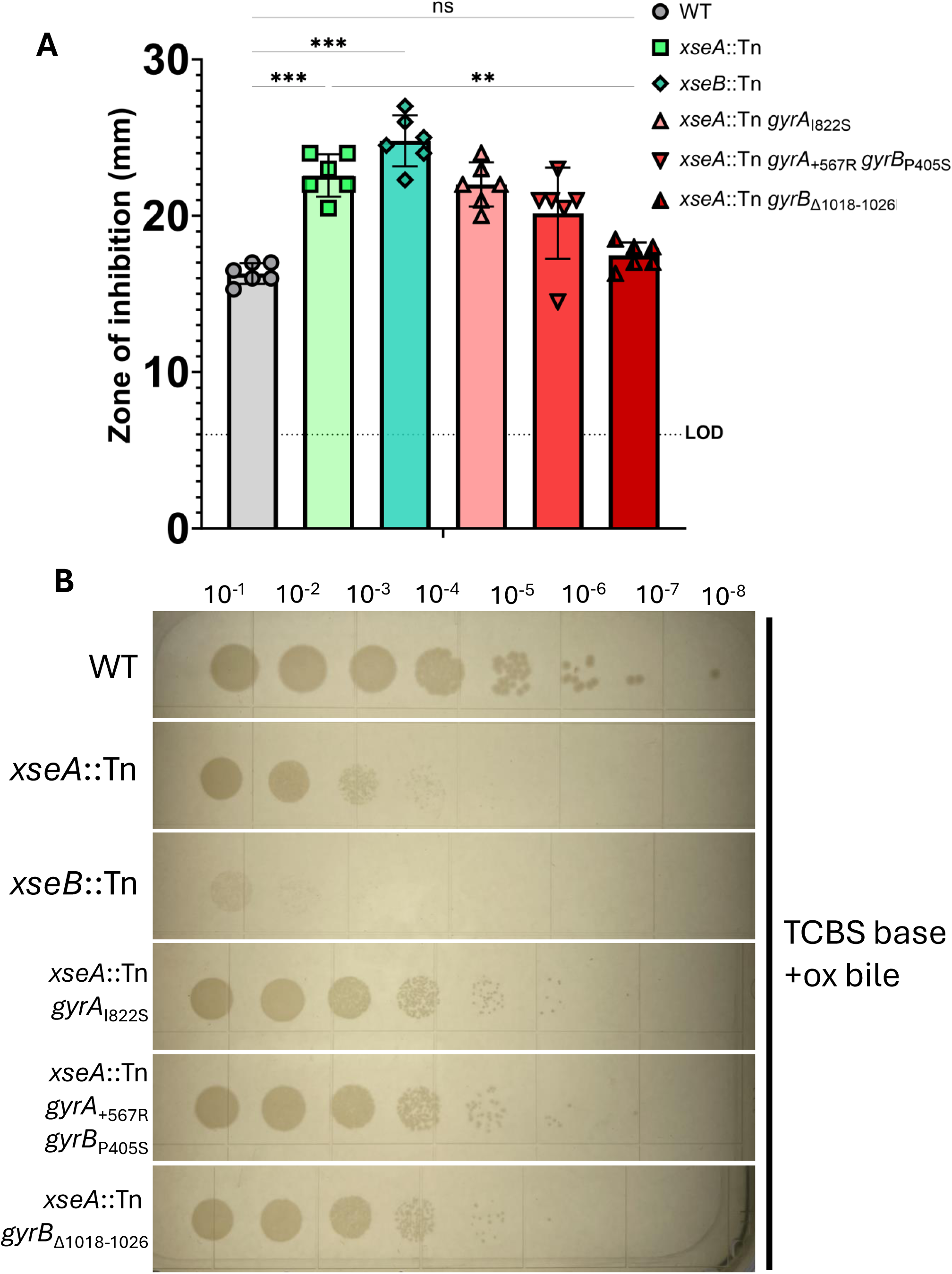
*exoVII* and *gyrAB* alter *V. cholerae* ciprofloxacin resistance. (A) WT, *exoVII* and *exoVII* suppressor mutants were assayed for ciprofloxacin sensitivity by disc diffusion with 5 ng of ciprofloxacin on LB agar. Results shown are from 6 biological replicate experiments. The dotted line at 6mm is labeled “LOD” for limit of detection as discs were 6 mm in diameter. Results were analyzed by one-way repeated measures ANOVA with Bonferroni’s multiple comparisons test. ns, P > 0.05; *, P ≤ 0.05; **, P ≤ 0.01; ***, P ≤ 0.001. (B) Serial dilution and spot plating of WT, *exoVII* and *exoVII* suppressor mutants over TCBS base + ox bile agar. Plates shown are representative of 3 biological replicate experiments.

Given that all three sets of DNA gyrase mutations suppress *xseA*::Tn TCBS toxicity but only one of the three confers ciprofloxacin resistance, we determined if these suppressor mutations restored growth on bile by plating serial dilutions of the suppressor mutants on TCBS base agar supplemented with only 0.5% (w/v) ox bile (Fig. 3B). While the *xseA::*Tn and *xseB::*Tn mutants demonstrate strong attenuation when plated on ox bile, all three suppressor mutants display significantly restored growth although not to the levels observed for WT *V. cholerae*. From these results we conclude that ox bile induces DNA gyrase-mediated DNA damage, similar to but distinct from ciprofloxacin, and the ExoVII null mutants fail to grow on TCBS because they are unable to repair this damage.

### Passaging *V. cholerae* on TCBS agar selects for ciprofloxacin resistance

We next hypothesized that if TCBS and ciprofloxacin inhibit the growth of the *exoVII* mutant via DNA gyrase toxicity, cells grown on TCBS should be more susceptible to inhibition by ciprofloxacin than cells grown on LB. We tested this using disc diffusion assays with the WT strain on LB agar and TCBS agar, and as predicted the zones of inhibition by ciprofloxacin of cells plated on TCBS agar was significantly greater than that of cells grown on LB agar (Fig 4A). This suggests that the additional damage caused by TCBS predisposes *V. cholerae* to ciprofloxacin toxicity. Next, because we found that the *gyrB*_Δ1018-1026_ suppressor mutation increased ciprofloxacin resistance of the *xseA*::Tn mutant, we tested if this *gyrB* mutations would confer resistance to ciprofloxacin in the WT strain. Our results demonstrated that the *gyrB*_Δ1018-1026_ strain produced a substantially smaller zone of inhibition than WT, suggesting that this mutant is resistant to ciprofloxacin even if *xseA* is functional (Fig. 4B).

**Figure 4:**
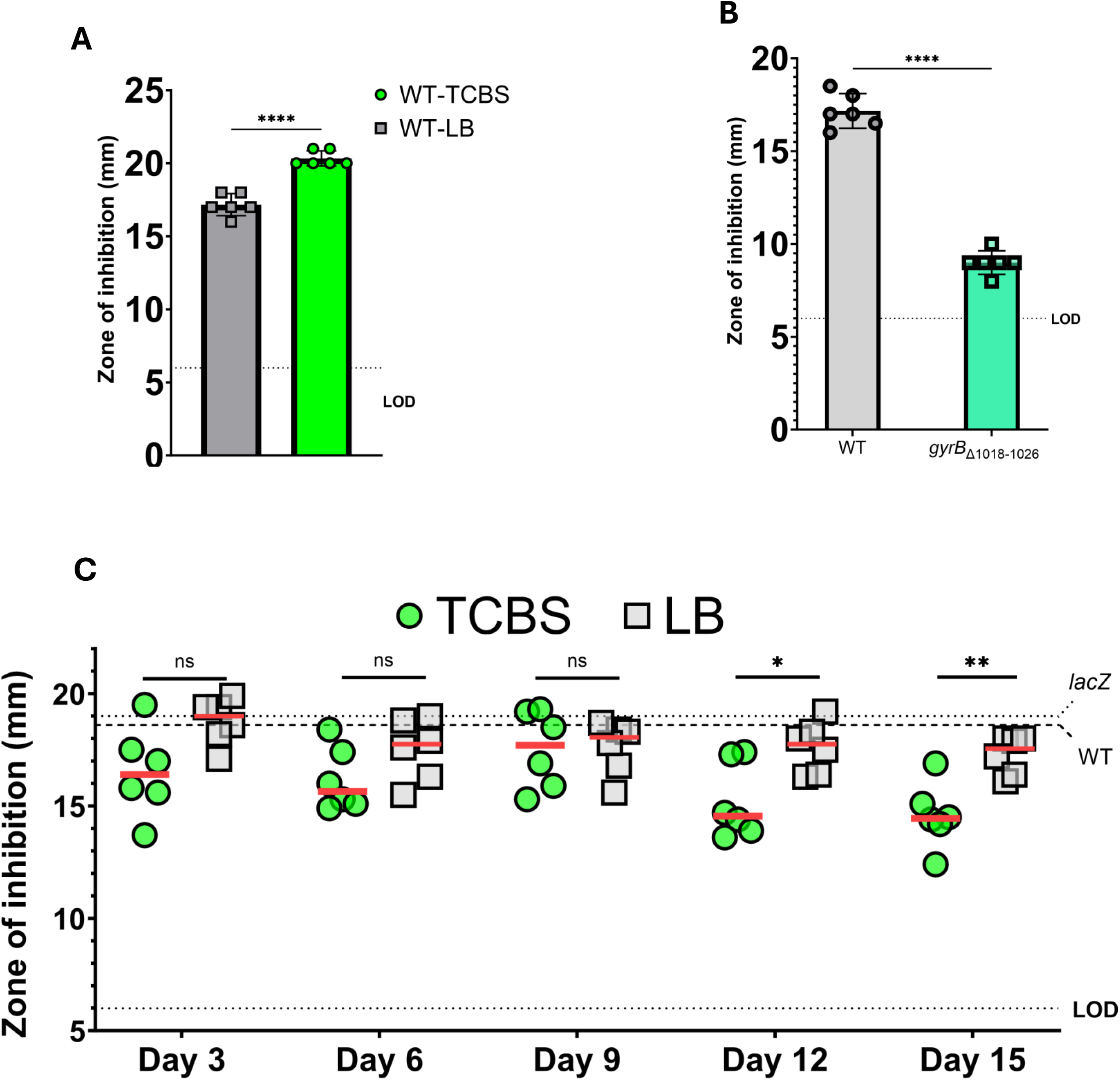
Passaging *V. cholerae* on TCBS agar selects for ciprofloxacin resistance. (A) WT *V. cholerae* was assayed for ciprofloxacin sensitivity by disc diffusion with 5ng of ciprofloxacin on LB or TCBS agar. Results shown are from 6 biological replicate experiments. Results were analyzed by unpaired T test using the Holm-Sidak method. P <0.0001. (B) WT *V. cholerae* and a strain carrying the *gyrB*_Δ1018-1026_ mutation were assayed for ciprofloxacin sensitivity by disc diffusion with 5ng of ciprofloxacin on LB agar. Results shown are from 6 biological replicate experiments. Results were analyzed by unpaired T test with Welch correction using the Holm-Sidak method. P <0.0001. (C) Comparison of ciprofloxacin sensitivity of 6 randomly-selected isolated *V. cholerae* (*lacZ* mutant) colonies following consecutive daily passages on either TCBS or LB agar (see methods). Isolates were assayed for ciprofloxacin sensitivity by disc diffusion with 5ng of ciprofloxacin on LB agar. The red lines correspond to the median ZOI among the 6 isolates. The dashed lines labeled “*lacZ*” and “WT” represent the average (n = 3) ZOIs for the unpassaged *lacZ* parent strain and unpassaged WT *V. cholerae*, respectively. Each data point represents the ZOI of a distinct isolate. Results were analyzed by Mann-Whitney two-tailed T test between the two lineages at each passage shown. Day 12, P = 0.026; Day 15, P = 0.0087. In A, B and C, the dotted line at 6mm is labeled “LOD” for limit of detection as discs were 6mm in diameter.

Our results thus far suggest that *V. cholerae* growing on TCBS experiences DNA gyrase mediated DNA damage similar to ciprofloxacin treatment. We therefore tested whether continuous growth of the WT strain on TCBS agar would impact ciprofloxacin resistance by serially passaging on either TCBS or LB agar daily for 15 days. Every 3 days we isolated six random clones per lineage to assess their sensitivity to ciprofloxacin by disc diffusion. On days 12 and 15 we observed an increasingly significant difference in the distributions of ZOIs between the TCBS and LB lineages (Fig. 4C). Notably, while the TCBS lineage clones varied widely in their sensitivity to ciprofloxacin at every passaging interval, the LB lineage clones were relatively consistent with the unpassaged Δ*lacZ* parent strain. This result therefore suggests that passaging on TCBS agar potentiates the evolution of ciprofloxacin resistant *V. cholerae*.

### Deployment of a randomly-barcoded TnSeq (RB-TnSeq) library identifies DNA repair mutants attenuated for growth on TCBS agar

Given that our results thus far suggest TCBS agar causes DNA damage and is selective for *V. cholerae* growth, we hypothesized that other DNA repair pathways would be required for growth on TCBS agar. To explore this hypothesis and more generally assess genes necessary for growth of *V. cholerae* on TCBS, we constructed a randomly barcoded transposon insertion site sequencing (RB-TnSeq) (23) mutant library in *V. cholerae*. Our library contains over 36,000 barcoded transposon insertion sites mapped to the *V. cholerae* C6706 genome, with at least one barcoded transposon mapped to 2833/3571 protein-coding genes (79.33%) (Table 2) (33).

**Table 2.** Barcoded mutant *V. cholerae* library summary.

| **Table 2.) Barcoded mutant *V. cholerae* library summary.** |  |
| --- | --- |
| Mutants collected | 77,678 |
| Mapped barcodes | 43,751 |
| Off-by-one barcodes masked | 7,361 (16.82%) |
| Usable, mapped barcodes | 36,691 (83.86%) |
| Unique insertion sites | 29,988 |
| Central 80% CDS insertions | 23,252 |
| Unique central insertion sites | 19,108 (82.18%) |
| Protein-coding genes in *V. cholerae* | 3,571 |
| With central insertions | 2,833 (79.33%) |

We first used our RB-TnSeq library in experiments to screen for mutants that failed to grow. These experiments were done in triplicate in TCBS broth cultures that contained all components of TCBS (labeled “TCBS complete”) to maintain library diversity. By sequencing and counting the barcodes present in the selected mutant pool and comparing those counts to those obtained from multiple uncultured library aliquots (see methods), we calculated scores for 2649 *V. cholerae* genes.

Our screening approach was validated as we measured highly negative mutant phenotypes for a multitude of genes that are known to be necessary for survival in the presence of bile (Fig 5A). For example, the outer membrane porins OmpU and OmpT, which are known to interact differently with bile, provide a useful test case for the screen (34). OmpT transport is blocked by deoxycholic acid, and as such does not contribute to bile resistance (11). The OmpU porin, on the other hand, is transcriptionally upregulated in response to bile by ToxR and has been long implicated in *V. cholerae* bile resistance (7, 9). Concordantly, while *ompT* had a nearly neutral score (−0.022) for growth in TCBS in our screens, *ompU* had a highly negative score (−6.546) (Fig. 5A). We also measured deleterious phenotypes for mutants in *tolC* (−7.417), *vexAB* (−2.314 and −2.276) and *vexCD* (−1.98 and −1.914) (Fig. 5A). VexAB and VexCD are Resistance Nodulation Division (RND) family efflux pumps that function in conjunction with the outer membrane pore TolC to expel bile acids from the cell and thus, are known to be involved in *V. cholerae* resistance to bile (35–38).

**Figure 5:**
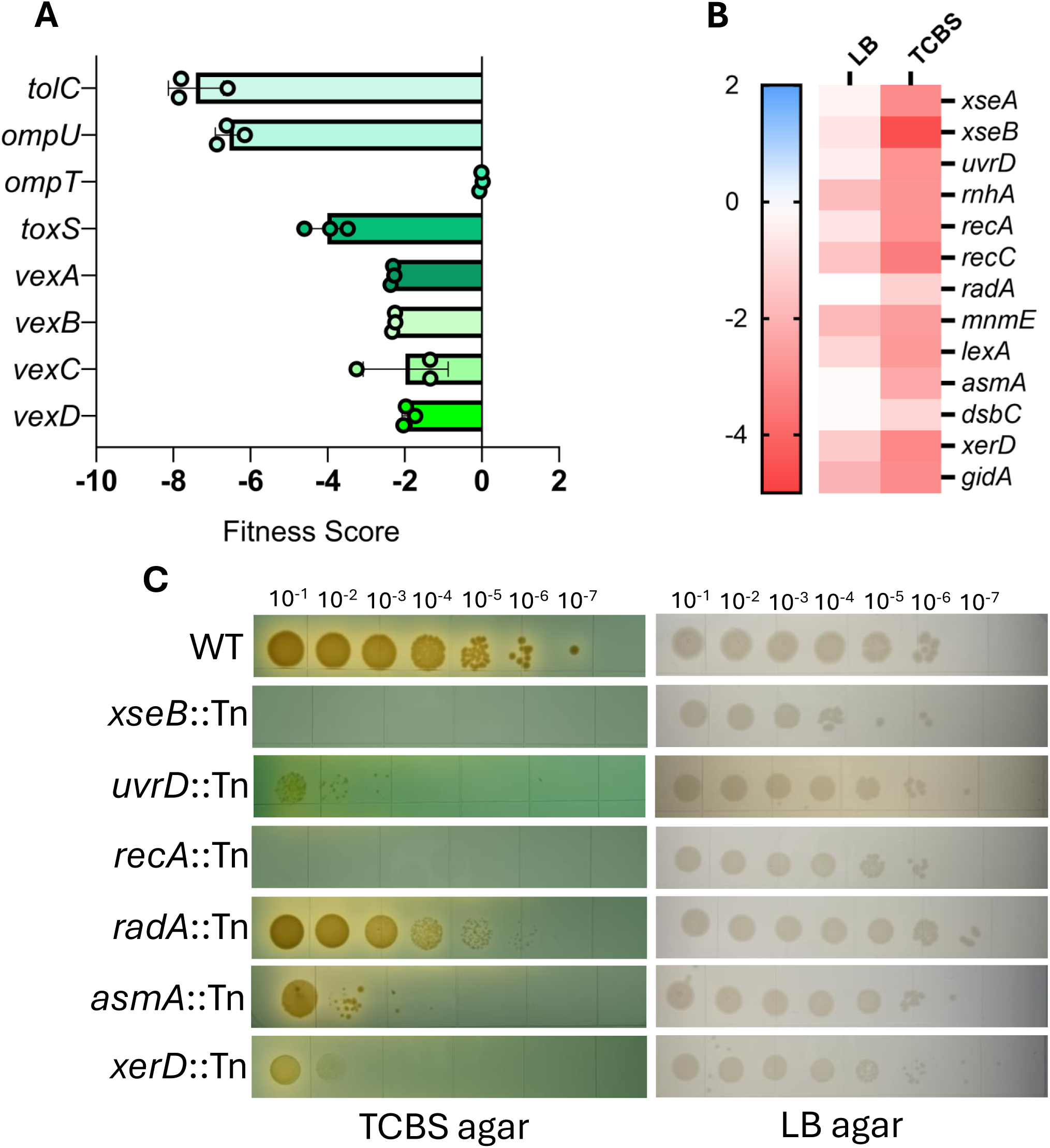
Deployment of an RB-TnSeq library identifies DNA repair mutants attenuated for growth on TCBS agar. (A) Mutant scores for genes known to be required for *V. cholerae* bile tolerance are shown. In each bar, each datapoint represents the gene score measured in 1 of 3 biological replicate experiments. A negative gene score indicates interruption of that gene by transposon insertion had a deleterious effect, while a positive score indicates a growth advantage. (B) DNA repair genes meeting quality thresholds (see methods) identified by gene ontology enrichment using ShinyGO 0.80 are shown. Gene scores are the average of 3 biological replicate experiments. (C) Serial dilution and spot plating of WT and DNA repair mutant strains over TCBS agar and LB agar. Plates shown are representative of 3 biological replicates per strain.

Given the success of our screen in identifying genes that could be predicted to be required for growth in TCBS, we next examined the rest of the genome. To identify genes that are specifically required for TCBS and do not exert general toxicity, we compared the datasets generated from growth in TCBS broth to a similar analysis of the library grown in LB using the same conditions. Genes were filtered for specific defects as described in the materials and methods. Filtering the data using these metrics yielded 111 genes that, when interrupted by Tn insertion, cause strong defects to *V. cholerae* grown in TCBS broth relative to LB (Fig. S1A). Gene ontology term enrichment analysis using ShinyGO (25) on these 111 genes revealed enrichment for genes involved in 4 major cellular processes: 1.) phospholipid transport, membrane organization and lipopolysaccharide (LPS) biosynthesis, 2.) phosphate ion transport, 3.) DNA replication and repair, and 4.) carbohydrate import and metabolism (Fig. S1B). Of note, we identified negative scores for many genes implicated in DNA repair (Fig. 5B). We retrieved several mutants in DNA repair genes from the ordered mutant library and demonstrated they had decreased growth on TCBS, but not LB, relative to the WT strain (Fig. 5C).

### RB-TnSeq reveals both component-dependent and independent growth defects for *V. cholerae* mutants in TCBS

Given that TCBS is a complex media with many components (Table 1), we wondered how each selective ingredient impacted the gene fitness scores observed upon selection with complete TCBS. We once again leveraged the high-throughput qualities of RB-TnSeq by repeating the TCBS (“complete” medium) screens in TCBS broth base (“base” medium) and TCBS broth base (“base+1”) media supplemented with either sodium citrate, ox bile, sodium cholate, ferric citrate, or sodium thiosulfate (Table 1).

Principal component analysis of each replicate dataset from both the TCBS complete, base, and base+1 experiments show strong clustering of replicate experiments (n=3). Interestingly, all the base+1 media datasets cluster together with the exception of base +ox bile (Fig 6A), demonstrating that the addition of ox bile applies a unique, strong selective pressure to the pool of mutants relative to the other selective components. Interestingly, TCBS complete media is distinct from every other dataset, implying that the multiple inhibitory components in TCBS complete apply a combinatorial selective pressure on *V. cholerae* growth that cannot be attributed to individual components themselves.

**Figure 6:**
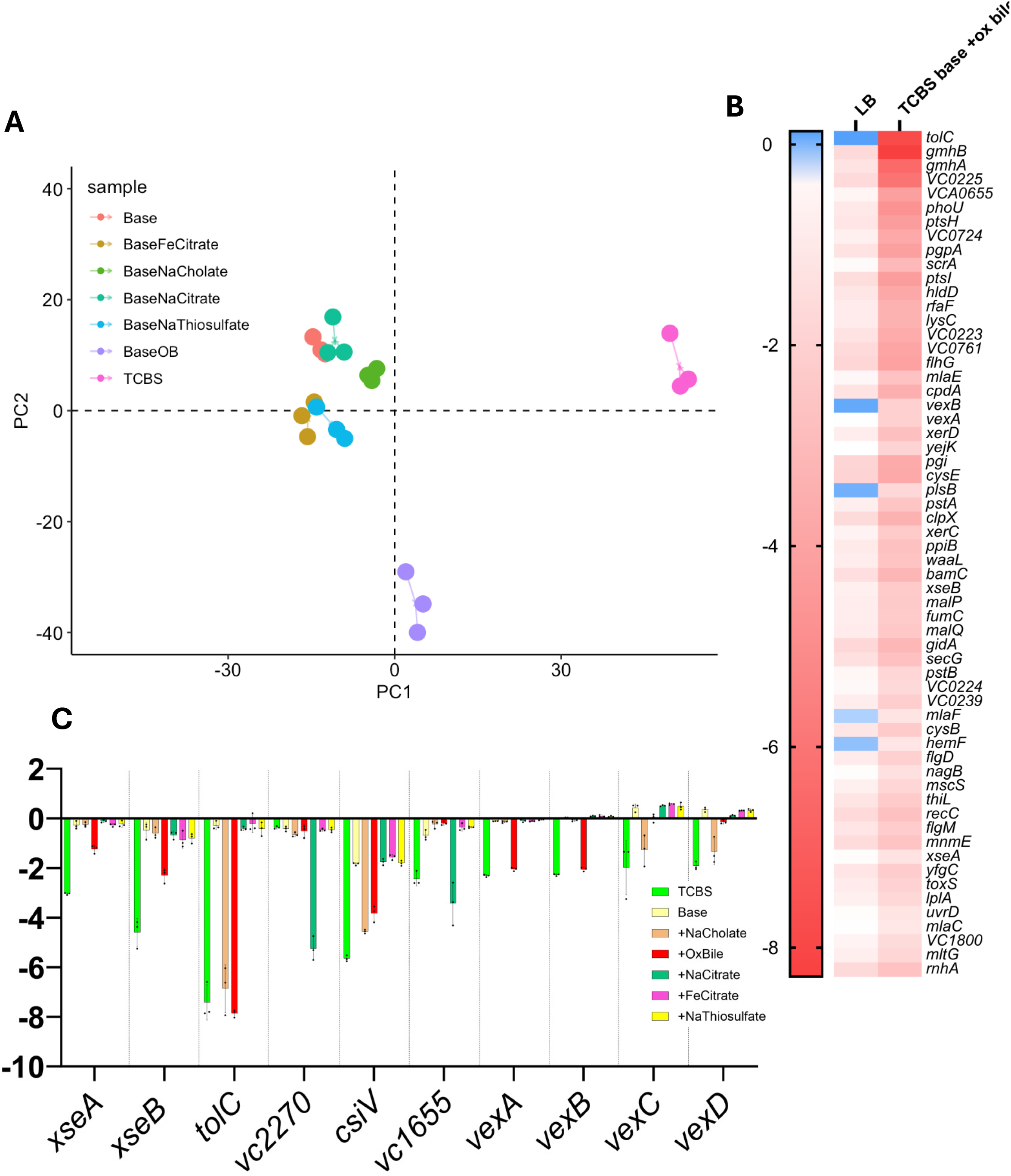
RB-TnSeq reveals both component-dependent and independent growth defects for *V. cholerae* mutants in TCBS. (A) Principal component analysis of TCBS complete and TCBS base (with and without individual selective components, Table 1) by gene scores. For each condition, individual experimental replicates are plotted in the same color and connected by lines. Plot was generated in R version 4.3.2 using the package vegan. (B) A wide array of mutants are attenuated for growth in TCBS base supplemented with ox bile. Genes shown in heatmap meet quality thresholds (see methods). Gene scores are the average of 3 biological replicate experiments. (C) Gene scores across media demonstrate component-dependent defects for *V. cholerae* mutants. Each dot in each bar represents the gene’s score in one of three biological replicate experiments. Dashed vertical lines were added to separate genes.

Given the observed clustering of the TCBS base +ox bile datasets relative to the rest of the base+1 experiments, we first examined the results of the TCBS base +ox bile dataset using the filtering metrics previously described. Filtering genes and scores by these metrics yielded a list of 60 genes that exhibited specific growth defects in TCBS base + ox bile (Fig. 6B). As expected, we had strong gene fitness scores for *xseA* and *xseB* in TCBS complete and TCBS base +ox bile, confirming our previous results (Fig. 6C). Notably, no other strongly negative scores were observed for *xseA* and *xseB* in any other TCBS base+1 condition, consistent with our results from Fig. 1B. This result provides confidence that we have captured genuine component-dependent fitness defects in our barcoded mutant *V. cholerae* strains.

We next searched for component specific defects by compiling all gene scores across the TCBS base+1 screens and sorted them by their standard deviation across conditions, revealing an assortment of genes with highly negative scores in only one or a few conditions tested (Fig. 6C). *tolC* mutants were highly attenuated in TCBS base supplemented with either ox bile or sodium cholate, but they did not have a fitness defect in sodium citrate, iron citrate, or sodium thiosulfate. Other genes were specifically important for growth in sodium cholate (*vexC* and *vexC*) or ox bile (*xseA*, *xseB*, *vexA*, and *vexB*).

Additional component-dependent fitness defects were observed for other genes. For example, transposon mutants in *vc2270* (riboflavin synthase alpha subunit, *ribC*) and *vc1655* (magnesium transporter, *mgtE*) did not impact growth in bile acids but showed reduced fitness the base +sodium citrate condition (Fig. 6C). It has been shown previously that *V. cholerae* lacking the peptidoglycan binding protein *csiV* is hypersensitive to the bile acid deoxycholate (39). Thus, we were not surprised to find *csiV* mutants, though selected against in all conditions tested, were most strongly inhibited by sodium cholate and ox bile. These results demonstrate that each selective component of TCBS requires a distinct set of genes to allow growth, supporting that TCBS complete media is the most selective (Fig. 1B) and distinct by PCA analysis (Fig. 6A).

## Discussion

Although TCBS has been used to isolate *V. cholerae* for decades, there is limited understanding of how this media is selective for *Vibrios* and the traits of *V. cholerae* that are critical for growth. Here, using a combination of forward and reverse genetics, we establish that *V. cholerae* experiences significant DNA damage when growing on TCBS and thus DNA repair is a key phenotype that promotes growth.

We serendipitously found that ExoVII is necessary for *V. cholerae* growth on TCBS agar, and that ox bile is largely responsible for the attenuation of *exoVII* mutants. Additionally, we found that suppression of the *exoVII* TCBS growth defect is through DNA gyrase mutations. Interestingly, some, but not all, of these DNA gyrase mutations also confer ciprofloxacin resistance. Previously, ExoVII had no well-described role outside of functional redundancy with ExoI and SbcCD exonucleases in recombinational repair (40) and ExoI and RecJ in methyl-directed mismatch repair (MMR) (41). Recently, ExoVII of *E. coli* was shown to be the exonuclease that is capable of excising quinolone-induced trapped DNA gyrase cleavage complexes (19). Together, this evidence suggests that the components of TCBS agar (most likely ox bile) cause ciprofloxacin like DNA damage to *V. cholerae*, explaining why *exoVII* mutant *V. cholerae* are inhibited by TCBS.

We can only speculate how the *gyrA* and *gyrB* mutations suppress the *exoVII* mutant’s sensitivity to ox bile. Even more interesting is the nature of the *gyrB*_Δ1018-1026_ mutation that confers suppression to *exoVII* mutants on TCBS and ciprofloxacin resistance, considering the most prevalent mechanism for evolving quinolone resistance in bacteria involves missense mutations in the quinolone resistance determining region (QRDR) of *gyrA* (amino acids 67-106 according to *E. coli* KL16 numbering) (42, 43). Because these residues allow targeting of quinolones to DNA gyrase/topoisomerase IV through a conserved water-metal ion bridge (44, 45), it is unlikely that any of the *gyrA* or *gyrB* mutations presented here alter the affinity of ciprofloxacin for DNA gyrase. This could indicate that the inhibition of *exoVII* mutant *V. cholerae* on TCBS occurs in a mechanism distinct from quinolones, though both insults result in lethality due to accumulation of trapped DNA gyrase cleavage complexes. Further mechanistic studies are required to disentangle the similarities and differences of how ox bile and ciprofloxacin cause DNA damage.

Besides the DNA gyrase mutations, one of the four suppressor mutants was found to have a frameshift mutation in *dinG* that likely results in a null phenotype. DinG is an ATP-dependent, structure specific helicase with 5’ → 3’ directionality (46). *In vitro* studies using a variety of substrates indicate that DinG is active on structures that mimic replication and homologous recombination intermediates (47). It has been suggested that DinG works semi-redundantly alongside helicases Rep and UvrD to permit DNA replication fork progression through transcribed regions by either displacing R-loops or dislodging RNA polymerase (48). It is currently unclear why a null mutation in *dinG* would suppress the toxicity of TCBS to the *exoVII* mutant. However, *dinG* was found to be upregulated in *E. coli* following treatment with the quinolone nalidixic acid (49). Additionally, *N. meningitidis dinG* mutants were found to be more sensitive to DSBs caused by mitomycin C than WT (50). We speculate that, in the absence of capable ExoVII, DinG may be overactive and thus is deleterious for *V. cholerae* cells grown on TCBS.

In this work we also confirm that, like in *E. coli* and *S. agalactiae*, *V. cholerae exoVII* mutants are hypersensitive to ciprofloxacin (19, 51), highlighting a common role for ExoVII in excising quinolone-induced trapped DNA gyrase cleavage complexes among these bacteria. In doing so we also report a role for ExoVII in surviving bile-mediated DNA damage. It is of particular interest how ox bile may mimic the DNA-damage caused by quinolones. An additional complication arises considering that ox bile as a poorly defined mixture of 9 individual bile acids, with different suppliers having varying abundances of each bile acid (52).

It is well understood that bile possesses antimicrobial properties, though our understanding of the exact mechanism(s) through which killing occurs is incomplete. A previous study found that the bile acids chenodeoxycholate and deoxycholate activate the SOS response in *E. coli*. Additionally, the study demonstrates that *E. coli* cells treated with these bile acids induce the SOS response and are killed similar to mitomycin C treatment (18). Other studies have suggested roles for base excision repair, SOS-induced DNA repair, and recombinational repair mediated by RecBCD in *S. enterica* for tolerating bile-induced DNA damage (13, 14). Moreover, evidence from both gram positive and negative bacteria suggests that resistance to bile is multifaceted. Several enteric bacteria express efflux pumps capable of expelling bile salts from the cell, thereby preventing extensive membrane damage caused by the detergent-like properties of bile. However, when membranes are disrupted and the cell’s permeability barrier is disturbed bile salts may enter cells and cause DNA damage, halting replication and leading to cell death in the absence of appropriate DNA repair mechanisms (12). TCBS is routinely used in laboratories to confirm the identity of *V. cholerae* strains. We demonstrate here that extended passage of *V. cholerae* on TCBS agar selects for ciprofloxacin resistant isolates and potentially against mutants that have defects in DNA repair machinery. Therefore, serial passaging on TCBS might be mutagenic and thus should be performed cautiously. We suggest against the practice of screening DNA repair mutant strains over TCBS for confirmation of *Vibrios* since some DNA repair mutants fail to grow completely and may accumulate further mutations when cultured over TCBS.

More importantly, bacterial pathogens can often evolve to have a mutator phenotype during infection. This is typically a result of deficiencies in MMR (53), and can allow bacteria to gain genetic diversity to allow rapid evolution to new environments. Hypermutators have been observed in enteric pathogens like *E. coli* and *S. enteritidis* (54). Moreover, increased propensity for mutation is likely to play a role in the long-term colonization of cystic fibrosis (CF) patients, from which hypermutable *Burkholderia pseudomallei* (55), *Haemophilus influenzae* (56) and *Pseudomonas aeruginosa*(57) have been isolated. For *V. cholerae,* such mutators are not historically thought to evolve during infection. However, a recent metagenomic study of cholera patients and their household contacts indicated the presence of hypermutable *V. cholerae* in clinical isolates from both symptomatic and asymptomatic individuals based on sequence analysis (58). Our results suggest an intriguing possibility that TCBS itself may be selecting against mutator strains that evolve *in vivo* leading their underrepresentation in clinical *V. cholerae* isolates.

## Materials and Methods

### Growth of *V. cholerae* strains

Unless otherwise noted, wild type (WT) and mutant *V. cholerae* strains were cultivated from frozen glycerol stocks on shaking incubators set to 37°C with shaking at 210rpm. Strains to be plated in any derivation of TCBS agar were grown in TCBS base broth (homebrew) using the wt/vol amounts listed in Table 1, while strains plated on LB agar were grown in LB broth. All strains are described in Table S2 while all primers are described in Table S3. When necessary, growth media was supplemented with kanamycin (100 μg/mL), chloramphenicol (10 μg/mL), and/or trimethoprim (10 μg/mL).

### Generation or retrieval of mutant *V. cholerae* strains

Strain TND0252, TND0252, SAD238 and SAD530 were generously gifted to us by Ankur Dalia, Indiana University. Strains CW2171, CW2172 and CW2173 were generated by using natural transformation (MuGENT) as previously described (20). Briefly, *tfox*, *recJ* and *xseA* genes were amplified from WT C6706 gDNA and then used to repair these three mutations individually in the TND0252 parent background by co-transformation with a Δ*vc1807::trimR* fragment. The Δ*vc1807::trimR* fragment was generated by using primers ABD346 and ABD347 to amplify it from strain SAD530. Strain JBG013 was made using primers JBG019 and CW2709 to amplify the mutant Δ*lacZ* allele from strain SAD238 (20). We then used MuGENT to transform WT *V. cholerae* with the Δ*lacZ* allele and Δ*vc1807::trimR*, yielding strain JBG013.

Unless otherwise noted, transposon mutant strains were retrieved from an ordered mutant library (28). *xseB*::Tn was made by amplifying the region surrounding the *xseB* locus with primers xseB_2.5kb_F and xseB_2.5kb_R (Table S3), and then using an *in vitro* Tn5-Kan^R^ transposon insertion kit (Biosearch Technologies Inc, TNP92110). Competent *V. cholerae* was prepared by growing strain NG001(harboring plasmid pMMB-*tfox*-*qstR*) in LB broth with chloramphenicol and 100 µg/ml isopropyl β-D-1-thiogalactopyranoside (IPTG) to maintain the plasmid and induce natural competence, respectively. Competent NG001 was diluted 1:4 in 0.5x instant ocean before tDNA was added. We then transformed WT C6706 with the *xseB*::Tn fragment as previously described (21) and confirmed mutation by sanger sequencing before curing the mutant of the competence-inducing plasmid.

### Curing strains of pMMB-*tfoX*-*qstR*

Sequence confirmed mutants were isolated and cultured in LB with the appropriate antibiotic overnight. The following day, the culture was struck over an LB plate to obtain isolated colonies. After incubation overnight, isolated colonies were selected and patch plated over LB + kanamycin agar and LB + chloramphenicol agar. Cured clones were identified as those that grow on LB + kanamycin agar but do not grow on LB + chloramphenicol agar.

### Spot plating of bacterial strains

Strains were grown as previously described overnight before being subcultured 1:100 in the same medium. Once reaching exponential growth strains were diluted in PBS to an OD_600_ of 0.5. Serial 10-fold dilutions were made and 2 μL were plated in spots and allowed to dry. Plates were then incubated at 37°C overnight.

### Generation of *xseA*::Tn suppressor mutants

*xseA*::Tn *V. cholerae* C6706 was retrieved from an ordered mutant library and cured of the competence-inducing plasmid (28). *xseA*::Tn was then cultured in LB overnight and 100 μL of the saturated culture was plated on TCBS agar. The plate was incubated at 37°C for 48 hours, after which isolated colonies were selected for follow up experiments. We extracted genomic DNA from the mutants using the Promega Wizard Genomic DNA Purification Kit (A1120). Genome sequencing of mutants was performed by SeqCoast Genomics in 150bp paired-end reads on an Illumina NextSeq 2000 and mutations were predicted using BreSeq (22).

Suppressor mutations were determined by comparing the BreSeq outputs for our evolved suppressor mutants against the BreSeq outputs generated by sequencing an assortment of repaired ordered library strains. We then amplified the *gyrA* (using primers gyrA_F and gyrA_R, Table S3) and/or *gyrB* (using primers gyrB_F and gyrB_R, Table S3) alleles from these mutants and transformed them into either *xseA*::Tn or WT parent strains using pMMB-*tfox-qstR* as described above. Mutations were confirmed by whole genome sequencing before strains were cured of the competence-inducing plasmid.

### Disc diffusion assays

Whatman filter paper was cut into discs using a hole puncher and sterilized by autoclaving. Strains were grown overnight and then back diluted 1:100 before being allowed to grow to exponential phase. Cells were then diluted to OD_600_=0.1 in PBS and 100 μL was plated on LB agar or TCBS agar. Discs were then placed with sterile forceps onto inoculated plates before being impregnated with different dosages of ciprofloxacin or a 0.1N HCl vehicle control. Plates were incubated at 37°C overnight and zones of inhibition were measured the following day.

### Serial passaging over TCBS agar

A single culture of Δ*lacZ V. cholerae* was grown overnight in LB+ trimethoprim. The following morning the culture was back diluted 1:100 before being allowed to grow to exponential phase. Cells were then diluted to OD_600_ to 0.5, and 100 μL of diluted culture was plated on either LB or TCBS agar. Plates were incubated at 37°C overnight. The following day, a sterile 10 μL loop was used to collect samples from bacterial lawns, which were then resuspended in 10 mL PBS. Samples were then diluted further in PBS to OD_600_ of 0.5 and 100 μL of the dilutions were replated on either LB or TCBS agar. Plates were again incubated at 37°C overnight. This process was repeated for 15 days, with passaging happening once per day per lineage (LB lineage and TCBS lineage). Every day 1 mL freezer stocks were made from the PBS resuspensions by mixing 750 μL with 250 μL 80% glycerol and storing at −80°C.

Every three days, the resuspensions were struck on LB + trimethoprim plates that were then incubated at 37°C overnight. The following morning, 6 isolated colonies per lineage were picked and cultured in 1mL LB broth at 37°C. Cultures were allowed to grow to exponential phase, back diluted to OD_600_ of 0.1, and 100 μL of each was plated over LB agar. We then assayed ciprofloxacin sensitivity using the disc diffusion assays described above.

### Construction and mapping of the RB-TnSeq mutant *V. cholerae* library

The *E. coli* APA752 donor strain harboring the pKMW3 transposon vector (kindly gifted to us by Aretha Fiebig, Michigan State University) was grown overnight in LB broth with 0.3 mM diaminopimelic acid (DAP) and kanamycin. We introduced pKMW3 into WT *V. cholerae* through conjugation. The APA752 culture and a WT *V. cholerae* culture were combined in a 9:1 ratio and before 100 μL was plated over LB with 0.3 mM DAP and incubated at 37°C overnight. The following day, bacteria were collected with sterile loops, resuspended in 50 mL LB, and pelleted by centrifugation. The supernatant was removed, and cells were washed in 15 mL LB to remove residual DAP. 5 mL of 80% glycerol was added, and the entire 20 mL volume was split up between 10×2 mL frozen stocks. Transposon mutant titer was measured by plating transconjugants over LB + kanamycin agar. Afterwards, 200 μL of thawed transconjugant stocks were plated over twenty-five 150×15mm petri dishes containing LB agar with kanamycin. Plates were incubated at 37°C for 48 hours.

Following incubation, three representative plates were selected, and colonies were counted to estimate the total mutant colonies collected between all 25 plates. For each plate, 3 mL of LB was added, and colonies were resuspended using sterile L-shaped spreaders. We collected as much of the 3 mL volume as possible from each plate and pooled it all in a single 50 mL conical. Afterwards, 30 mL of the collected cells were added to a flask containing 270 mL LB with kanamycin, and the mutants were incubated at 37°C with 210 rpm shaking for 1 hour. Glycerol was then added to a 20% final concentration and hundreds of 100 μL library aliquots were frozen for single use, as well as a few dozen 1mL aliquots to facilitate efficient genomic DNA extraction. We extracted genomic DNA from the mutants using the Promega Wizard Genomic DNA Purification Kit.

For library mapping, we followed the strategy of Wetmore and colleagues (23) with minor modifications to the PCR enrichment of barcode-containing insertion junctions. The modifications are described in detail elsewhere (24). Briefly, 5 μg of library genomic DNA was sheared with a Covaris M220 ultrasonicator to produce ∼300 bp fragments. Genomic DNA was then electrophoresed on a 1% TBE gel, stained with SYBR Gold (Invitrogen, S11494), and gel extracted using the Zymoclean Gel DNA Recovery Kit (Zymo Research, D4001) to select for fragments between 150-500 bp. We then performed end repair, A-tailing and adaptor ligation with the NEBNext Ultra II DNA Library Prep Kit for Illumina (E7645S) following the manufacturer’s recommended protocol using custom adaptors made by annealing oligos Mod2_TruSeq and Mod2_TS_Univ (Table S3). Adaptor-ligated DNA was cleaned without size selection using NEBNext Sample Purification Beads (0.9x) and eluted in 0.1x TE buffer following the manufacturer’s instructions.

DNA fragments containing the transposons were enriched in a 2-step nested PCR strategy using modified primer sequences based on the original primer sequences used by Wetmore et al (23). This modified enrichment strategy is described by Fiebig and colleagues (24). In the first PCR we used the forward primer TS_pHimar+4_F and the reverse primer TS_R (Table S3). Cycling parameters were 98°C for 2 min, 10x (98°C, 30sec; 70°C, 20sec; 72°C, 20sec), 72°C for 5min, and a 4°C hold. After the first PCR, products were cleaned again using NEBNext Sample Purification beads (0.9x) and eluted in 0.1x TE buffer. 15 μL of the cleaned PCR product was used as template in the second transposon enriching PCR. In the second PCR we used the primers P7_MOD_TS_index3 and P5_TS_F (Table S3) to add Illumina P5 and P7 sequences and a 6-bp i7 index. Cycling parameters were 98°C for 3 min, 10x (98°C, 20sec; 70°C, 10sec; 72°C, 20sec), 72°C for 5min, and a 4°C hold. Following the second PCR, products were again cleaned using NEBNext Sample Purification beads (0.9x) and eluted in 0.1x TE buffer. Reagents for both PCR reactions were supplied in the NEBNext Ultra II DNA Library Prep Kit for Illumina. For both reactions we used the volumes of reagents as outlined by the kit manual. Prepared reads were submitted to SeqCoast Genomics (Portsmouth, NH) for sequencing on an Illumina NextSeq using a 300-cycle NextSeq P1 reagent kit (Illumina 20050264). The sequencing run was supplemented with 25% phiX DNA to aid in clustering. Locations of reliable transposon insertions were aligned and mapped to the *V. cholerae* C6706 genome (GenBank accession numbers CP064350 and CP064351) using custom Perl scripts written and described by Wetmore et al (23) (available at https://bitbucket.org/berkeleylab/feba/src/master/). Mapping statistics are provided in Table 2.

### Competition of RB-TnSeq *V. cholerae* mutants in liquid media

For each replicate RB-TnSeq experiment in TCBS broth (including TCBS with or without a selective component), we thawed a single 100 μL aliquot of the RB-TnSeq library on ice before being added to 100 mL of experimental liquid medium in 250 mL unbaffled Erlenmeyer flasks. Initial OD_600_ values were obtained before cultures were incubated for 5 hours at 37°C with shaking. OD_600_ values were taken hourly. After 5 hours, cultures were collected in 50 mL conical tubes and pelleted. Supernatants were removed and genomic DNA was extracted from pellets as described above. The comparison experiment performed in LB broth was done almost exactly as described above, except that the cultures were incubated until they had reached OD_600_ of ∼1.0 (4 hours and 10 minutes)

The amplification and sequencing of barcodes was performed following the approach described by Wetmore et al (23). 100-200 ng of genomic DNA from each experiment sample was used as template to amplify mutant barcodes by PCR. Barcodes were amplified in 50 μL reaction volumes using Q5 DNA polymerase (New England Biolabs) using 1x Q5 reaction buffer, 1x High GC enhancer, 1.0 units Q5 polymerase, 0.2 mM dNTP and 0.4 μM of each primer. The forward primer was BarSeq_P1, while each reaction used a uniquely indexed reverse primer BarSeq_P2_ITXXX, where “XXX” corresponds to the index number as used by Wetmore and colleagues (Table S3).

Reaction conditions were as follows: 98°C for 4 min, 25x (98°C, 30sec; 55°C, 30sec; 72°C, 30sec), 72°C for 5min, and a 4°C hold. Following, samples were run on a 2% TAE agarose gel to confirm the presence of PCR products. 5 μL of each PCR was then pooled, and the pool of PCR products was cleaned using a DNA Clean & Concentrator kit (Zymo Research, D4003).

Barcodes were then sequenced by SeqCenter (Pittsburgh, PA) on a single lane of a NovaSeq X Plus flowcell in 2×150-cycle configuration. Barcodes were supplemented with 50% phiX DNA to aid in clustering and loaded at 110 pM. Reads were trimmed to 75 bp before analysis to eliminate unnecessary Illumina adaptors, leaving the barcode sequence. Barcode sequences in each sample were counted and the fitness of each mutant strain was calculated as the normalized log2 difference in barcode counts in the treated sample versus replicate reference samples using the custom scripts developed by Wetmore et al. For our reference samples, we extracted genomic DNA from four individual uncultured 100 μL library aliquots, amplified the barcodes and sequenced them as described above. Gene scores were calculated based on the weighted average of strain scores for mutants with insertions in the central 80% coding region of the gene, again using the scripts developed by Wetmore et al. (available at https://bitbucket.org/berkeleylab/feba/src/master/).

### Gene Ontology enrichment using ShinyGO

We wanted to identify genes that were specifically important for TCBS growth and not standard rich media. To identify these genes, we made the following comparison of fitness scores in TCBS to those in LB. Having calculated gene scores based on differences in barcode abundance relative to an uncultured library sample, we then filtered our analysis to exclude genes that have a score in TCBS or TCBS base +ox bile ≤-1 (mutant must be attenuated for growth in the experimental condition), ≥-2 in LB (to exclude those mutants with general fitness defects in rich media), and a difference in scores between the two conditions (experimental condition score – LB score) ≤-1. Using this thresholding strategy, we identified 111 genes necessary for growth in TCBS broth and 60 genes necessary for growth in TCBS base + ox bile broth, relative to their necessity for growth in LB.

We then performed gene ontology enrichment by comparing the gene IDs against annotations present in the *V. cholerae* N16961 genome (taxonomy ID 243277) using ShinyGO 0.80 (25). To eliminate error in the false discovery rate produced by ShinyGO, we submitted a background list of genes that included the gene IDs for all 2649 scored across the RB-TnSeq experiments. This way, enrichment of genes meeting our filtering criteria isn’t biased by the absence of genes we were unable to assign scores to in our RB-TnSeq experiments. ShinyGO options were as follows: FDR cutoff = 0.05, # pathways to show = 10, pathway size minimum = 10. For clarity, we then performed some manual binning of pathways based on redundancy between pathway names and the genes belonging to them (for example, pathways “DNA replication, and DNA repair”, “DNA metabolic process, and chromosome segregation” and “DNA repair, and DNA-dependent ATPase activity” contain many of the same genes, so we group them together more broadly under “DNA repair”).

### Statistical analysis and data visualization

All statistical analyses were performed in GraphPad Prism version 10.1.1 Principal component analysis plot was made in R version 4.3.2 (26) using the vegan package (27) version 2.6-6.1.

## ACKNOWLEDGEMENTS

This research was supported and funded by NIH grants GM139537 and AI158433 to C.M.W. We thank Jasper Gomez for constructing strain JBG013. We also thank Aretha Fiebig for sharing the strain and expertise necessary for generating the *V. cholerae* RB-TnSeq library and Ankur Dalia for sharing strains.

---

**<u>Figure S1: *V. cholerae* mutants attenuated for growth in TCBS broth</u>**. (A) All genes shown meet score thresholds as described in Methods. Gene scores shown are the average of 3 biological replicate experiments. (B) Genes from S1 that fit into a common cellular process are grouped by black labels. Gene scores shown are the average of 3 biological replicate experiments.

